# The *Streptococcus* phage protein paratox is an intrinsically disordered protein

**DOI:** 10.1101/2024.01.14.575580

**Authors:** Iman Asakereh, Nicole R. Rutbeek, Manvir Singh, David Davidson, Gerd Prehna, Mazdak Khajehpour

**Affiliations:** Department of Chemistry, University of Manitoba; Department of Microbiology, University of Manitoba

**Keywords:** Paratox, Prx, ComR, *Streptococcus*, quorum sensing, bacteriophage, intrinsically disordered protein, circular dichroism, NMR, protein folding

## Abstract

The bacteriophage protein paratox blocks quorum sensing in its streptococcal host by directly binding the signal receptor and transcription factor ComR. This reduces ability of *Streptococcus* to uptake environmental DNA and protects phage DNA from damage by recombination. Past work characterizing the paratox:ComR molecular interaction revealed that paratox adopts a well-ordered globular fold when bound to ComR. However, solution-state biophysical measurements suggested that paratox may be conformationally dynamic. To address this discrepancy, we investigated the stability and dynamic properties of paratox in solution using circular dichroism, nuclear magnetic resonance, and several fluorescence-based protein folding assays. Our work shows that under dilute buffer conditions paratox is intrinsically disordered. We also show that the addition of kosmotropic salts or protein stabilizing osmolytes induces paratox folding. However, only the addition of ComR was able to induce paratox to adopt its previously characterized globular fold. Furthermore, as we can induce different paratox folding-states we characterize Prx folding thermodynamics and folding kinetics using stopped flow measurements. Based upon the kinetic results, paratox is a highly dynamic protein in dilute solution, folding and refolding within the 10 ms timescale. Overall, our results demonstrate that the streptococcal phage protein paratox is an intrinsically disordered protein in a two-state equilibrium with a solute-stabilized folded form. Furthermore, the solute-stabilized paratox fold is likely the predominant form of paratox in a solute-crowded bacterial cell. Finally, our work suggests that Prx binds and inhibits ComR, and thus quorum sensing in *Streptococcus*, by a combination of conformational selection and induced-fit binding mechanisms.

## INTRODUCTION

*Streptococcus pyogenes* or Group A Streptococcus (GAS) is a bacterial pathogen that is implicated in several human diseases^1^. GAS can cause mild infections such as impetigo^2^ and pharyngitis^3^ or deadly conditions such as rheumatic fever^4^ and toxic shock syndrome^5^. Central to the pathogenicity of GAS is the presence of bacteriophage encoded toxin genes^6^. These toxin genes are found within the bacteriophage genome and are actively spread among different strains and species of *Streptococcus* through direct phage infection^7^. In fact, bacteriophage infection is a primary driving force of clonal diversity in GAS^8-9^.

After infection, lysogenic GAS bacteriophages incorporate their DNA into the GAS genome and persist as a stable prophage^10^. The prophage often encodes a deadly toxin gene near the 3’ end of the prophage. These toxin genes include the superantigen SpeA which is responsible for scarlet fever and toxic shock syndrome^11-12^. Adjacent to the toxin gene is the *prx* gene that encodes the protein paratox (Prx)^13-14^. Furthermore, the toxin gene and *prx* are one genetic cassette that remains intact when the GAS phage exits the lysogenic cycle and self-excises from the GAS genome^13^. The exact purpose of this linkage remains unclear, but current work strongly suggests that the linkage with *prx* is to maintain the genetic integrity of the toxin^14-15^.

Supporting this hypothesis is the observation that Prx inhibits natural competence in *S. pyogenes*^14^. Natural competence (natural transformation) in Streptococcus is a quorum-sensing regulated process in which bacteria control the expression of a number of genes that encode the machinery for both the acquisition and incorporation of DNA^16^. In GAS, natural competence is regulated by the ComRS quorum-sensing pathway^17-19^. ComR is a quorum-sensing receptor and a transcription factor that is part of the RRNPP protein family^17^. ComS is a secreted peptide pheromone that is processed into a mature form termed XIP (SigX inducing peptide)^19^. Upon binding to ComR, XIP induces a large conformational change in ComR that results dimerization and the ability to bind to the promoter regions of *comS* (XIP) and *sigX*^20-21^. SigX then leads to the expression of late genes required for natural competence^22-23^. The expression of Prx is also induced by SigX, resulting in its own ComRS quorum-sensing regulated expression^19^. Prx then acts as a negative regulator of the system by binding to the DNA-binding-domain (DBD) of ComR directly blocking interaction with DNA and inhibiting the expression of competence genes^14-15^.

Prx is a small 60 amino acid protein that has been shown to adopt a globular fold in X-ray crystal structures alone^14^ or when bound to ComR^15^. However, previous biophysical characterization of Prx using small-angle X-ray scattering coupled to size exclusion chromatography (SEC-SAXS) shows that Prx occupies a larger volume than what is predicted from the folded crystal structure^15^, even though analytical ultracentrifuge (AUC) data clearly demonstrates that the protein is a monomer in solution^14^. The disagreement between the Prx crystal structures and its behavior in solution suggests that Prx is likely dynamic and adopts conformations significantly different from those captured in the crystal structures.

To probe this hypothesis, we have investigated the folding thermodynamics of Prx using circular dichroism (CD), fluorescence, and protein NMR. Our results show that under dilute and physiological solvent conditions, Prx has all the hallmarks of an intrinsically disordered protein^24^. Furthermore, the addition of salts and molecular crowders induce Prx to adopt a folded form. However, only the presence of its binding partner ComR induces Prx to adopt the previously characterized globular fold. We demonstrate that the folding of Prx before interaction with ComR can be properly described by a two-state thermodynamic model^25-27^ and propose that under physiological conditions Prx should be assumed to be at dynamic equilibrium between intrinsically disordered unfolded states and a solute-stabilized fold. This suggests that conformational selection^26, 28^ plays a critical role in the binding of Prx to ComR, and thus the inhibition of both quorum sensing and natural competence in *Streptococcus pyogenes*.

## RESULTS

### Structure of Prx in solution

As our past results suggest that the X-ray crystal structures of Prx are not representative of its fold solution^14-15^, we proceeded to assay the structure of Prx in solution. However, all previous experiments used a Prx expression construct that included a non-cleavable C-terminal 6His-tag. It is not uncommon for cloning artifacts such as a 6His-tag to affect protein stability^29^ or even stabilize protein conformations as crystal packing artifacts^20, 30-31^. In fact, the Prx crystal dimer (PDBid: 6CKA) is stabilized by the C-terminal 6His-tag of a symmetry mate^14^. Given this, we proceeded to create a new Prx expression construct without a 6His-tag. Instead, Prx was cloned with a cleavable N-terminal GST-tag. As shown in Figure 1A, the new Prx construct is readily purifiable and binds to its known biological partner ComR. Additionally, our Prx ortholog of study from MGAS315 contains no tryptophan residue impeding the ability to assay its fold by fluorescence. To address this problem, we made a Prx point variant substituting in tryptophan at phenylalanine residue 31 (PrxF31W). This residue was chosen as it is partially buried in the X-ray crystal structure^14^. The variant PrxF31W was still able to bind ComR, demonstrating that this mutation did not inhibit the biochemical function of Prx (Figure 1B). Moreover, this shows that PrxF31W can still adopt the fold observed in the X-ray crystal structures that is necessary to bind ComR^15^.

**Figure 1.**
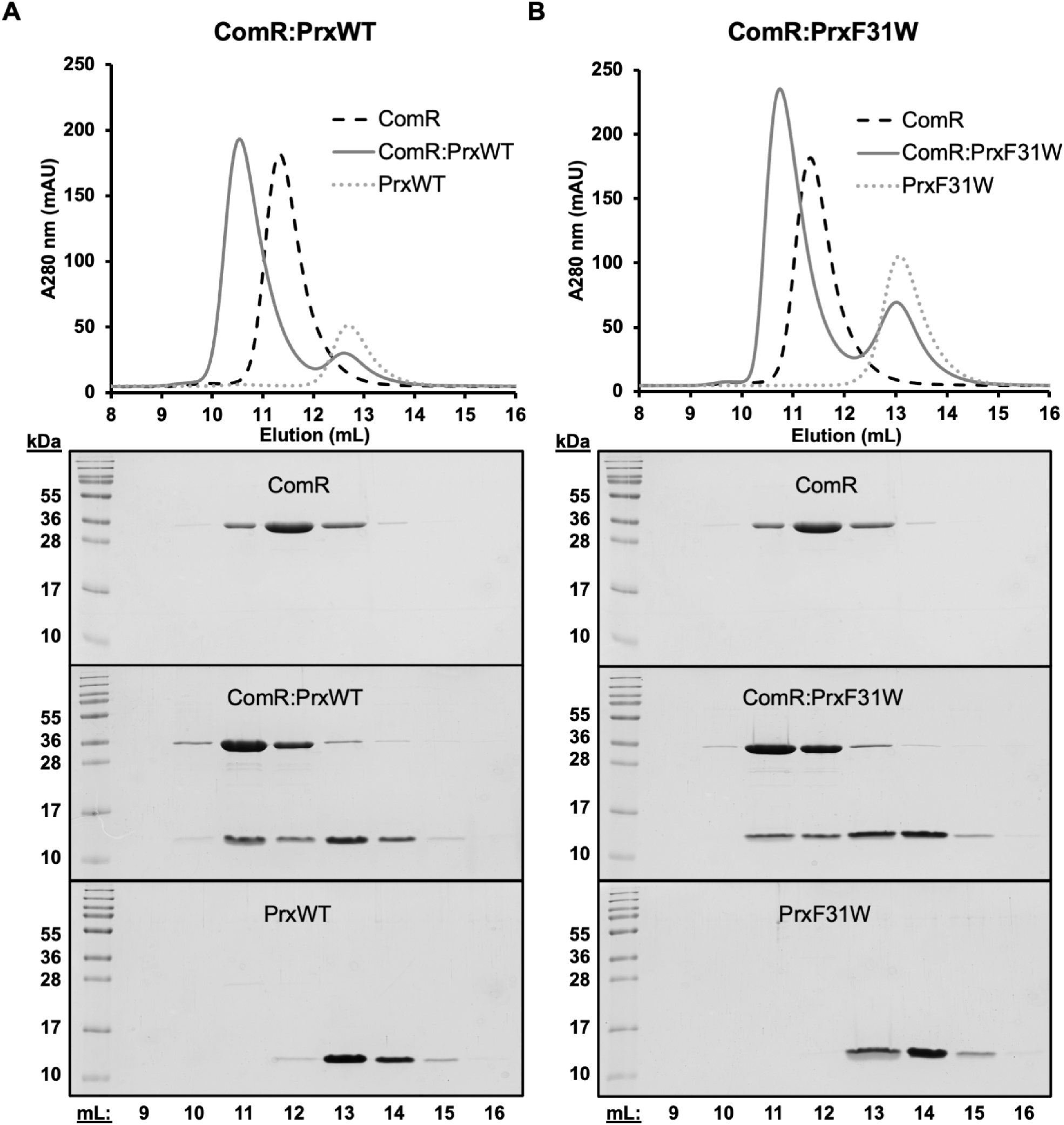
Purification of wild-type Prx and a PrxF31W variant. Size exclusion chromatography was performed with ComR:Prx complexes at a 1:1.5 molar ratio with **(A)** ComR and wild-type Prx and **(B)** ComR and PrxF31W. The upper panels for each show the SEC elution profile of the complexes overlayed over the elution profiles of ComR and PrxWT or PrxF31W. The lower panels show SDS-PAGE gels of the SEC eluted proteins or complexes, with each well corresponding to the elution volume of the SEC trace above.

We next proceeded to measure the structure of tag-less Prx in solution by circular dichroism (CD). Figure 2A demonstrates how the secondary structure of paratox is affected by salt concentration. It can be clearly seen that under low salt concentrations Prx lacks significant secondary structure, while the addition of KF induces folding. The spectral transition exhibits an isosbestic point around 205 nm, which is consistent with a two-state model for the folding process. We also measured the CD spectrum of the His-tagged Prx construct (Figure 2B). Our measurements agree with past CD data^15^ and indicate that the presence of the poly-histidine tag induces secondary structure in Prx.

**Figure 2.**
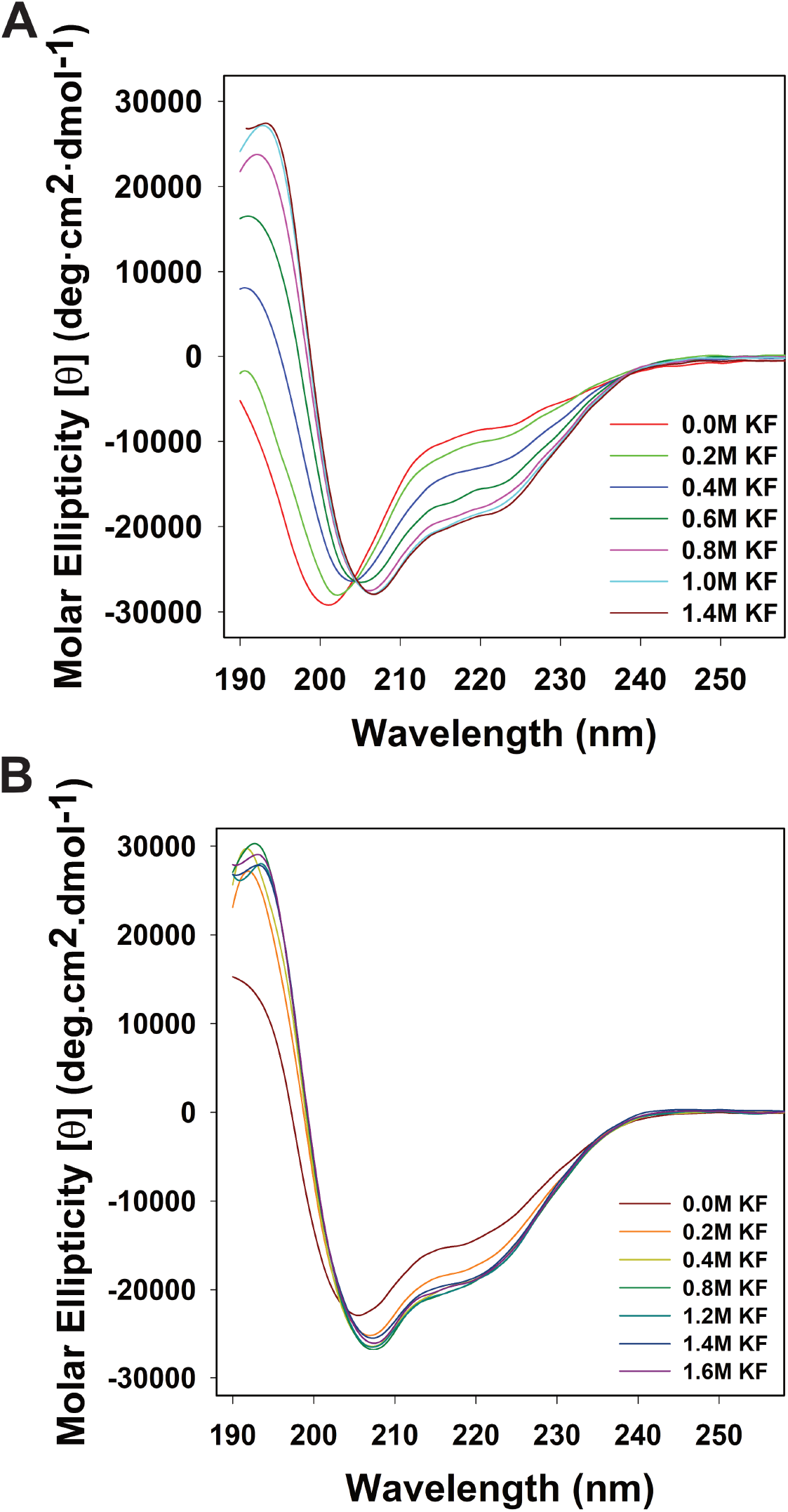
Prx is unfolded in dilute salt conditions. **(A)** Circular dichroism spectra of the Prx secondary structure as perturbed by the addition of potassium fluoride. All measurements were performed in triplicate at room temperature in 20mM Phosphate buffer (pH 7.0) with a final protein concentration of 20uM. **B)** Circular dichroism spectra of His-tagged Prx secondary structure as perturbed by the addition of potassium fluoride. All measurements were performed in triplicate at room temperature in 20mM Phosphate buffer (pH 7.0) with a final protein concentration of 20uM

As the CD spectrum demonstrated that the structure of Prx was unfolded at physiological buffer conditions, we then used NMR to gain further insight into the solution-state fold of Prx. Figure 3 depicts several ^1^H-^15^N HSQC spectra of ^15^N labeled Prx measured in various conditions. Figure 3A shows tag-less Prx in denaturing conditions (6.66 M Urea) and in buffer at pH 7. It can be clearly seen that in the absence of high-salt or crowding agents, the ^1^H-^15^N HSQC dispersion pattern of Prx at pH 7 is similar to that measured under denaturing conditions. Specifically, all backbone chemical shifts show low dispersion and are clustered between 8.0 ppm and 8.6 ppm in the hydrogen dimension. This ^1^H-^15^N HSQC pattern is diagnostic of a protein that lacks secondary structure and is disordered^32-33^. In contrast, the addition of unlabeled ComR DNA-binding domain (DBD) to ^15^N-labeled Prx results in a well resolved ^1^H-^15^N HSQC indicative of an ordered and globular protein fold (Figure 3C). Given the nano-molar affinity of the Prx:DBD interaction^15^ and the well-resolved spectra, we conclude that the HSQC in Figure 3B represents the Prx fold observed in the X-ray crystal structures.

**Figure 3:**
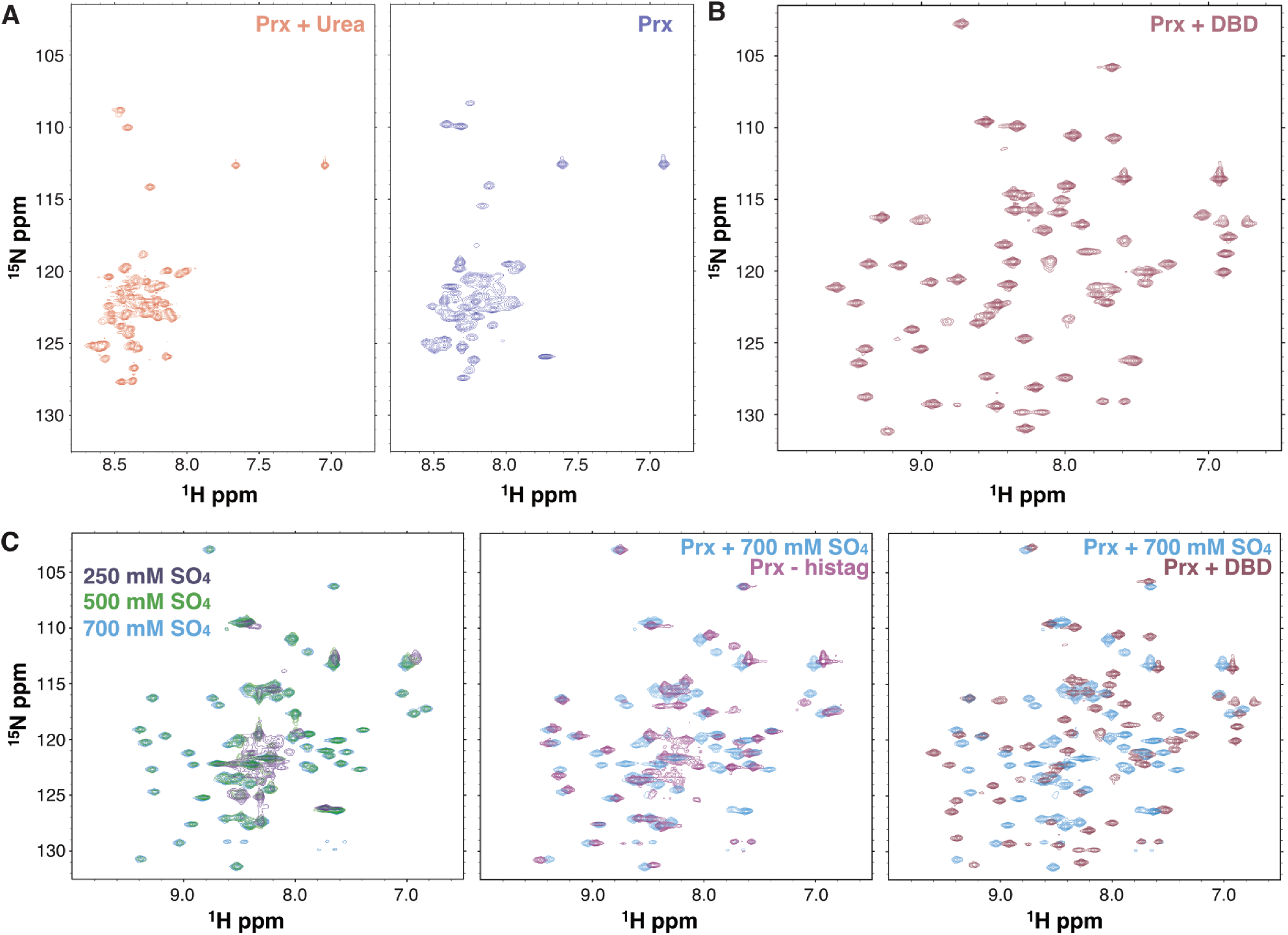
Paratox adopts disordered and ordered globular conformations. **(A)** ^1^H-^15^N HSQC spectra of ^15^N Prx dissolved in 6.66M Urea (left) and at pH 7.0 and low salt (right). **(B)** ^1^H-^15^N HSQC spectrum of ^15^N labeled Prx bound to unlabeled ComR DBD. **(C)** ^1^H-^15^N HSQC spectra of ^15^N Prx with increasing Na_2_SO_4_ concentrations (left). Overlay of a ^15^N Prx + 700 mM Na_2_SO_4_ spectrum with a spectrum of ^15^N Prx containing a 6His-tag (middle) or with a spectrum of ^15^N labeled Prx bound to unlabeled ComR DBD (right). In each panel the spectra are labeled by color.

Our CD measurements also showed that the structure of Prx was easily influenced by salt concentration. As such, we have also used NMR to probe the effects of salt upon the structure of Prx (Figure 3C). It can be clearly seen that the addition of sodium sulfate leads to an increase in peak resolution, inducing an ^1^H-^15^N HSQC pattern characteristic of a folded protein (Figure 3C, left). If we compare the ^1^H-^15^N HSQC of Prx in 700 mM sodium sulfate to that of the Prx 6His-tag in buffer at pH 7, we see that they are closely related with several overlapping peaks (Figure 3C, middle). This shows that the salt-induced fold of Prx is similar to the 6His-tag induced fold of Prx in agreement with our CD data (Figure 2B). However, comparing the ^1^H-^15^N HSQC spectrum of the salt-induced folded state with that of DBD-bound Prx (Figure 3C, right) shows that these Prx structures are not identical. This is even considering possible differences in Prx chemical shifts due to interaction with the ComR DBD. Taken together, our CD and NMR data demonstrate that Prx is an intrinsically disordered protein that can adopt a solute stabilized fold and ultimately a well-ordered rigid globular domain when bound to its biological interaction partner ComR. Furthermore, the salt and His-tag induced Prx folds are similar to each other but are different structures than those reported in the X-ray crystal structures.

### Folding properties of paratox

Given that Prx is dynamic and can adopt multiple different folds, we used the PrxF31W variant as a spectroscopic reporter to monitor protein folding. First, the KF induced folding of the F31W variant was also compared with that of wild type using CD (Figure 4A, B). The salt-induced folding profiles of wild-type and PrxF31W can be observed to be very similar, with the minor caveat that the substitution of phenylalanine by tryptophan may have slightly favored the folding of the protein. Next, the fluorescence properties of PrxF31W were tested in dilute buffer, high-salt, and denaturing conditions (Figure 5A). When dissolved in pH 7.5 tris buffer, the protein emission spectrum exhibits a peak around 354 nm consistent with that of a highly solvent-exposed tryptophan^34^. The addition of urea causes an additional red-shift in the protein spectrum, resulting in a fluorescence emission peak around 358 nm. In contrast, the addition of KF causes a blue shift in the protein spectrum, resulting in a measured emission peak around 350 nm. Thus, this data illustrates that the folding/unfolding of PrxF31W between the disordered and salt-stabilized form can be probed through monitoring its protein emission peak position. If the protein unfolds through a two-state mechanism, the observed maximum emission wavelength of the protein *λ*_*obs*_ will depend upon *x*_*F*_, the mole fraction of folded protein via the equation *λ*_*obs*_ = *x*_*F*_*λ*_*F*_ + (1 – *x*_*F*_)λ_*U*_, where *λ*_*F*_ and *λ*_*U*_ are the maximum emission wavelengths of the folded and unfolded states of the protein respectively^35^.

**Figure 4:**
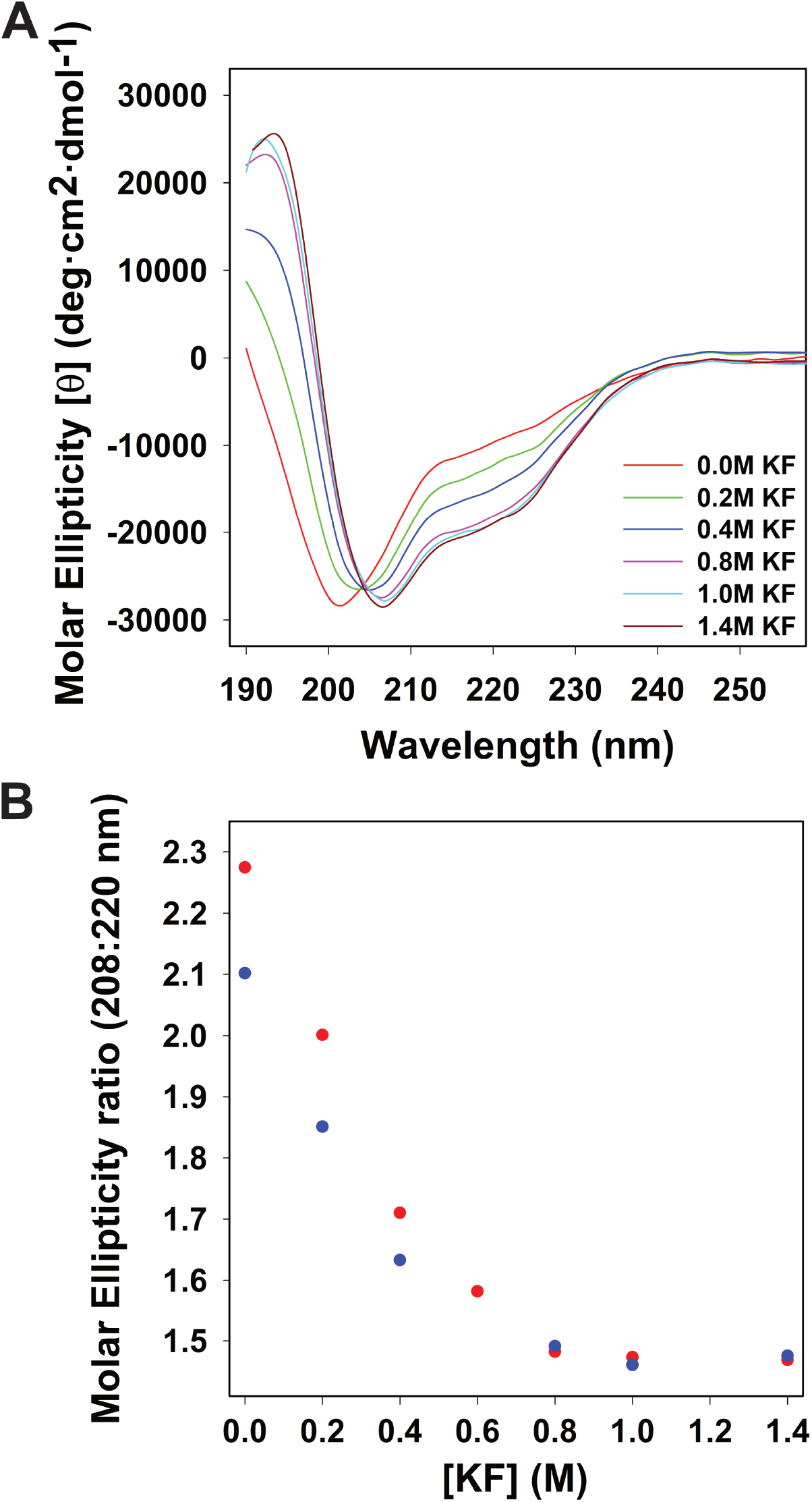
The Prx F31W variant has a comparable folding mechanism to wild-type Prx. **(A)** Circular dichroism spectra of PrxF31W secondary structure as perturbed by the addition of potassium fluoride. All measurements were performed in triplicate at room temperature in 20mM Phosphate buffer (pH 7.0) with a final protein concentration of 20uM. **(B)** Comparing the KF induced changes in helicity of wild-type paratox (red) and the paratox F31W variant (blue), as described by the 208:220 molar ellipticity ratios.

**Figure 5.**
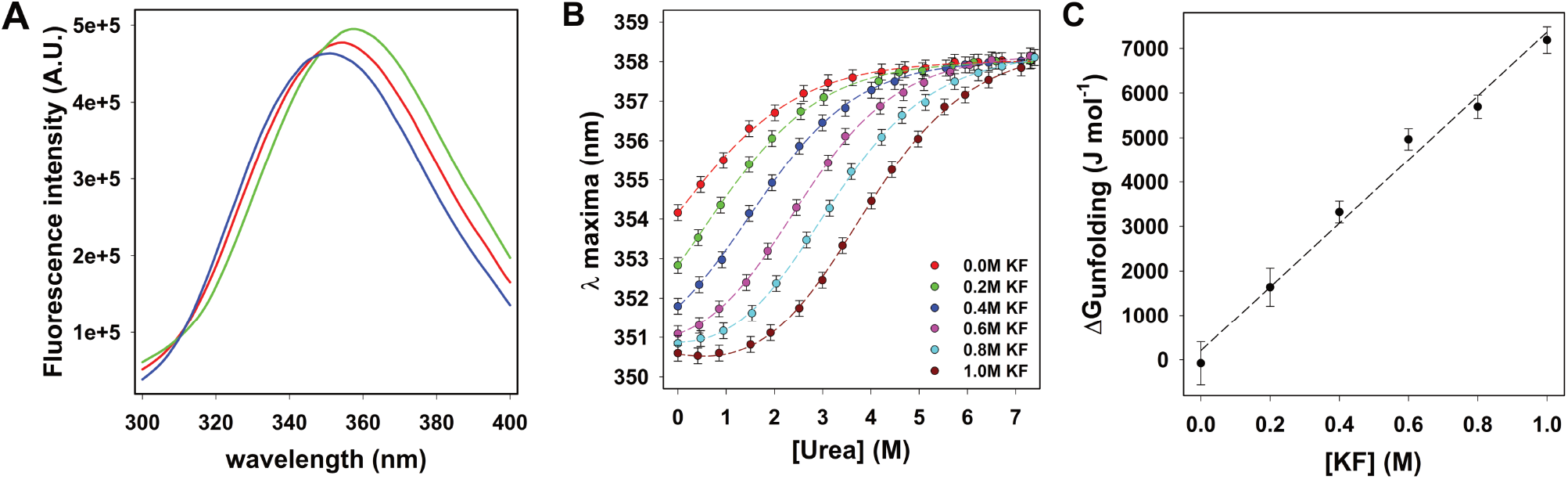
Fluorometric determination of Prx folding states. **(A)** Steady-state fluorescence spectrum of PrxF31W collected in 25mM Tris buffer (pH 7.5) buffer (red line), in 25mM Tris buffer (pH 7.5) containing 1.0 M KF (blue line) and in 25mM Tris buffer (pH 7.5) containing 7.1M Urea (green line). All measurements were performed at room 293 K with a final protein concentration of 5 μM. The samples were excited at 280 nm, with excitation and emission slits set to 2 nm bandpass. **(B)** Urea-induced unfolding profiles for PrxF31W measured in various KF solutions obtained by monitoring the *λ*_maxima_ of the protein fluorescence peak. All measurements were performed at 293 K, in 25mM Tris buffer (pH 7.5), with a final protein concentration of 5 μM. The samples were excited at 280 nM and the emission peak was collected from 300 to 450 nm, with both excitation and emission slits set to 2 nM bandpass. The *λ*_maxima_ was obtained by fitting each emission peak to a Weibull 5-parameter equation^59^, the error bars representing the uncertainty in *λ*_maxima_ obtained from fitting the data. The solid lines represent the best global fit of the data to Equation-1, sharing the parameters *λ*_U_ (357.6 ± 0.2 nm; the emission maximum of the unfolded protein) and *λ*_F_ (350.2 ± 0.1; the emission maximum of the folded protein). **(C)** Plotting the dependence of 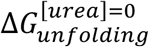obtained from the data in Table 1 upon KF concentration, the dashed line represents the best linear fit to the data.

The urea-induced denaturation profiles of Prx have been obtained by plotting the protein emission peak maxima as a function of urea concentration and different solvent buffer conditions containing varying amounts of KF (Figure 5B). As shown, the stability of the folded form of Prx increases with KF addition. This is clearly represented by the urea concentration associated with the folding transition midpoint. From these profiles the standard free energy of folding can be calculated from Eq-1^36^, assuming that Prx folds via a two-state mechanism:

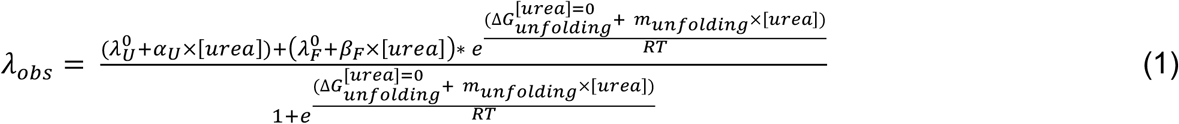

**Table 1.** Unfolding thermodynamic parameters of PrxF31W measured in various KF solutions. Parameters were obtained from the best global fit of data from Figure 4B to Eq-1, sharing λ_U_ = 357.6 ± 0.2 nm and λ_F_ = 350.2 ± 0.1nm, through non-linear least square repression analysis.

| [KF] (M) | $\frac{\Delta G_{unfolding}^{[urea]=0}}{RT}$ | $\frac{m_{unfolding}}{RT}$ | R<sup>2</sup> |
| --- | --- | --- | --- |
| 0.0 | -0.03 ± 0.04 | -0.8 ± 0.2 | 0.9988 |
| 0.2 | 0.67 ± 0.12 | -0.8 ± 0.1 | 0.9991 |
| 0.4 | 1.36 ± 0.09 | -0.91 ± 0.04 | 0.9994 |
| 0.6 | 2.30 ± 0.11 | -0.89 ± 0.02 | 0.9997 |
| 0.8 | 2.34 ± 0.11 | -0.89 ± 0.02 | 0.9997 |
| 1.0 | 2.95 ± 0.12 | -0.87 ± 0.04 | 0.9998 |

In this equation, 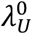 is the emission maximum wavelength of unfolded Prx in a given buffer in the absence of urea, 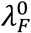 is the emission maximum wavelength of folded Prx in a given buffer in the absence of urea, *λ*_*obs*_ is the emission maximum of paratox measured at a given concentration of urea, *α*_*U*_ and *β*_*F*_ linearly correct for the effect of urea upon the spectrum of folded and unfolded 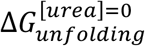 is the standard free energy of unfolding measured in the absence of urea, and *m*_*unfolding*_ is the urea denaturation “m-value” as defined by Pace^37^. The data in Figure 5B have been fitted to Eq-1, sharing the parameters 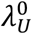 and 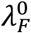 for all data sets. The resulting fitting parameters are tabulated in Table 1. The coefficients of determination (r^2^) for all data fits are greater than 0.99, suggesting that the folding of Prx is well-described by a two-state model. The “wellness” of these fits suggests that the presence of KF has small effect on the emission properties of folded and unfolded Prx, allowing the maximum emission wavelengths of folded and unfolded Prx in pH 7 tris buffer (25 mM) to be estimated: 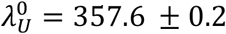 and 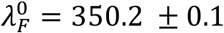. Additionally, the standard folding free energy of Prx in non-crowded dilute conditions 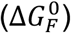 can be determined by plotting the values of 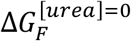 as a function of KF concentration (Figure 5B). As expected, these data are fairly well-correlated linearly (r^2^ ≈ 0.988) (Figure 5C); allowing for an estimate of: 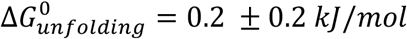. Prx is therefore clearly exists in a dynamic equilibrium between the folded and unfolded states under dilute conditions. The salt-induced folding mechanism of Prx was also studied with stopped-flow in buffers containing 1M and 0.5 M KF to observe the kinetics of Prx state transitions. Lower concentrations of salt were not studied by stopped-flow, because it is impossible to fully fold the protein at low salt concentrations. Figure 6 depicts typical stopped-flow traces of the unfolding/unfolding of Prx in buffer containing varying amounts of urea. Figure 6A represents PrxF31W folding traces and Figure 6B shows PrxP31W unfolding traces. The folding/unfolding fluorescent traces of Prx are well described by single-exponential kinetics as described by equations:

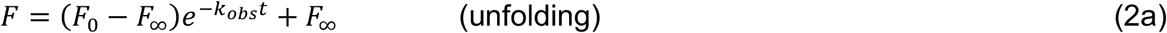

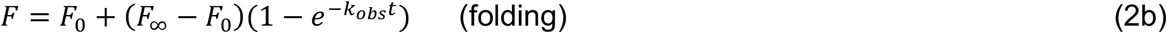

In these equations, *t* is time, *F*_0_ is the fluorescence intensity measured at *t* = 0, *F* _*∞*_ is the fluorescence intensity extrapolated to *t* = ∞, and *k*_*obs*_ is the apparent rate constant which for a two-state folding transition is equal to the sum of *k*_*folding*_ and *k*_*unfolding*_, the folding and unfolding rate constants. For all measured traces, the fitting parameters of Equations 2a and 2b are given in the supplementary data.

**Figure 6.**
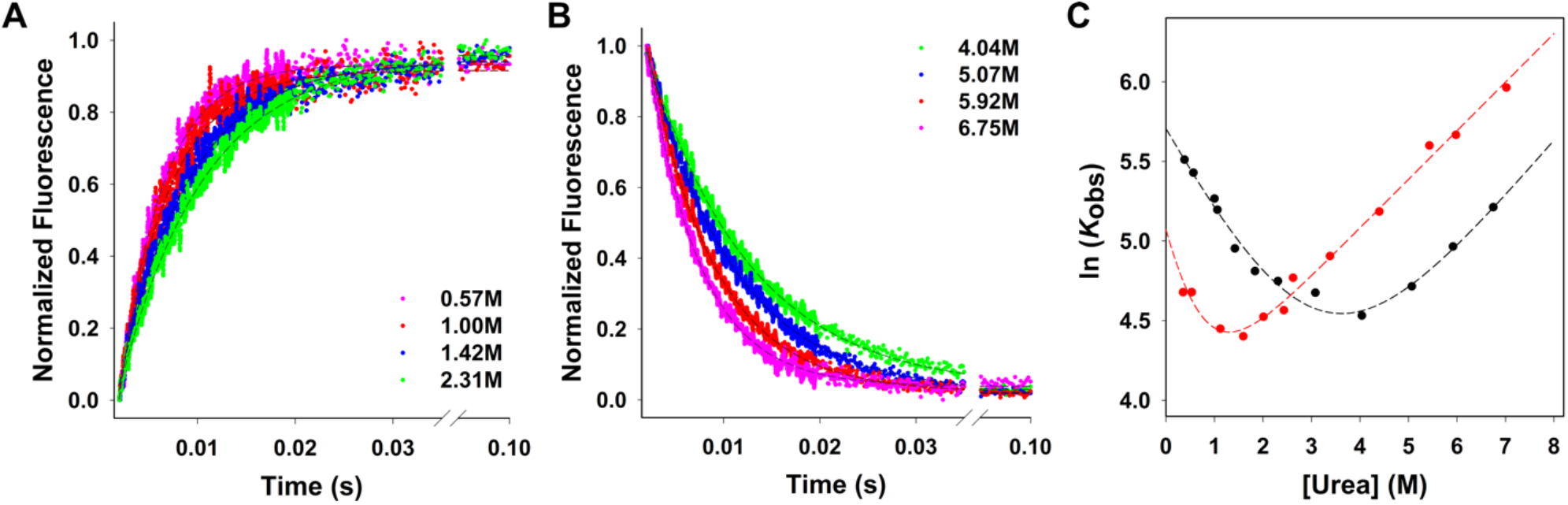
refolding and unfolding kinetics of PrxF31W shows a two-state system. Typical stopped-flow kinetic traces of PrxF31W unfolding and refolding measured in 25mM Tris buffered at pH 7.5, containing 1.0 M KF at 20 °C. **(A)** Prx refolding traces (fluorescence increase) measured in the following urea concentrations are depicted: 0.57M red, 1.00 M green, 1.42 M blue, 2.31 M pink. **(B)** Prx unfolding traces (fluorescence decrease) measured in the following urea concentrations: 4.04 M dark red, 5.07 M dark green, 5.92 M dark blue, 6.75 M dark pink. The change in protein fluorescence is monitored at 330 nm and is normalized to 1 for reasons of clarity. The solid and dashed lines represent the best mono-exponential fits (Equations 2a and 2b) to the obtained data. **(C)** Chevron plots depicting the observed folding and unfolding relaxation rate constants as a function of urea concentration, shown for 1.0M KF (black) and 0.5M KF (red). All measurements were performed in 25mM Tris buffered at pH 7.5, fitting errors are smaller than the circle diameters. For each KF concentration, the solid curves represent the best fit of the data to Equation-3.

We have further investigated the salt-induced folding of Prx through constructing a chevron plot^38^ from our obtained *k*_*obs*_ values as seen in Figure 6C. The dashed line represents the best fit to the equation:

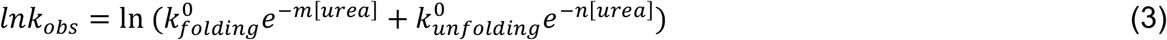

In which 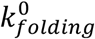 and 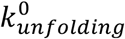 represent the folding and unfolding rate constants of the Prx in the absence of urea, while *m* and *n* are constants. The lack of any appreciable rollover observed in the chevron diagrams^39^ and the fact that their minimum points are consistent with the inflection points of the denaturation curves of Figure 5B, strongly suggests that the folding of Prx into the salt-stabilized fold follows a two-state mechanism.

From the chevron plots the folding and unfolding rate constants in 1 M KF 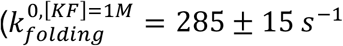 and 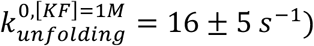 and in 0.5 M KF 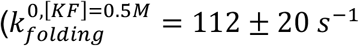 and 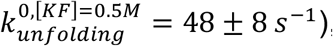, can be obtained. If the unfolding free energy of activation is extrapolated to 0 M KF, we can estimate the folding and unfolding rate constants in dilute pH 7 buffer to be: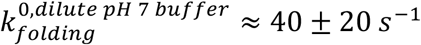 and 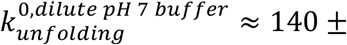 90 *s*^−1^ ; thus, indicating that the folded form of Prx in dilute buffer is dynamic and has lifetime significantly shorter than 10 ms, which is consistent with the disordered ^1^H-^15^N HSQC spectrum the protein exhibits in Figure 3.

### Salt and crowder effects upon the folding of paratox

The presence of salt can clearly affect the folding of Prx in solution. However, it is unclear if these effects are mediated through electrostatic screening or Hofmeister salting-out effects^40^. Figure 7A shows the effects of different salts upon the folding of Prx. Clearly, the salt induced folding of Prx exhibits strong ion-specificity, suggesting that Hofmeister effects are perhaps the dominant contributors to the process. Hofmeister effects on the protein folding standard free energy Δ*G*_*unfolding*_ are best expressed^40^ by the equation: 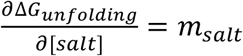. This allows us to modify the Eq-1 for this two-state protein to:

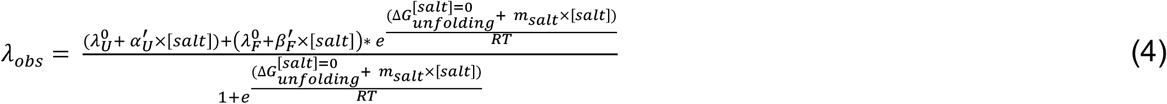

In this equation, 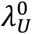 is the emission maximum wavelength of unfolded Prx in a given buffer in the absence of salt,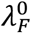 is the emission maximum wavelength of folded Prx in a given buffer in the absence of salt, 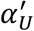 and 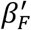 linearly correct for the effect of salt upon the spectrum of folded and unfolded Prx, 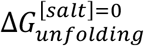 is the Prx standard free energy of unfolding measured in pure buffer in the absence of salt and *m*_*salt*_ is the “m-value” characterizing Hofmeister effects as defined by Record^40^. The data in Figure 7A have been globally fitted to Eq-4, sharing the parameter 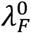 and constraining 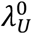 to be equal to 357.6 nm for all data sets. The resulting fitting parameters “*m*_*salt*_” and 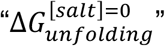 are tabulated in Table 2. The salt-induced folding data of Prx are well-corelated with Eq-4 (the coefficients of determination (r^2^) for all data sets are greater than 0.99). The absence of any appreciable nonlinear salt concentration dependence of Δ*G*_,*unfolding*_ emphasizes that electrostatic screening plays a minimal role in the salt-induced folding of paratox^40-42^.

**Table 2.** Salt induced folding parameters of parameters of PrxF31W measured in various saline solutions. Parameters were obtained from the best global fit of data from Figure 6A to Eq-4 and Figure 6B to Eq-4a, sharing λ_U_ = 357.6 ± 0.2 nm and λ_F_ = 350.2 ± 0.1nm, through non-linear least square repression analysis.

| Co-solute | $m/RT$ | $\Delta G_F^{[co-solute]=0}$<br>(kJ/mol) | R<sup>2</sup> |
| --- | --- | --- | --- |
| NaCl | 2.25 ± 0.03 | - 0.1 ± 0.2 | 0.9989 |
| NH<sub>4</sub>CL | 1.15 ± 0.14 | - 0.1 ± 0.2 | 0.9984 |
| LiCl | 1.09 ± 0.02 | - 0.2 ± 0.2 | 0.9986 |
| RbCl | 2.18 ± 0.04 | - 0.2 ± 0.2 | 0.9995 |
| KCL | 2.06 ± 0.04 | - 0.1 ± 0.2 | 0.9984 |
| KF | 3.79 ± 0.08 | -0.1 ± 0.2 | 0.9994 |
| Na<sub>2</sub>SO<sub>4</sub> | 7.59 ± 0.07 | -0.1 ± 0.2 | 0.9997 |
| xylitol | 1.33 ± 0.05 | -0.2 ± 0.2 | 0.9990 |
| proline | 1.30 ± 0.90 | - 0.2 ± 0.2 | 0.9946 |
| TMAO | 2.26 ± 0.05 | - 0.2 ± 0.2 | 0.9979 |

**Figure 7.**
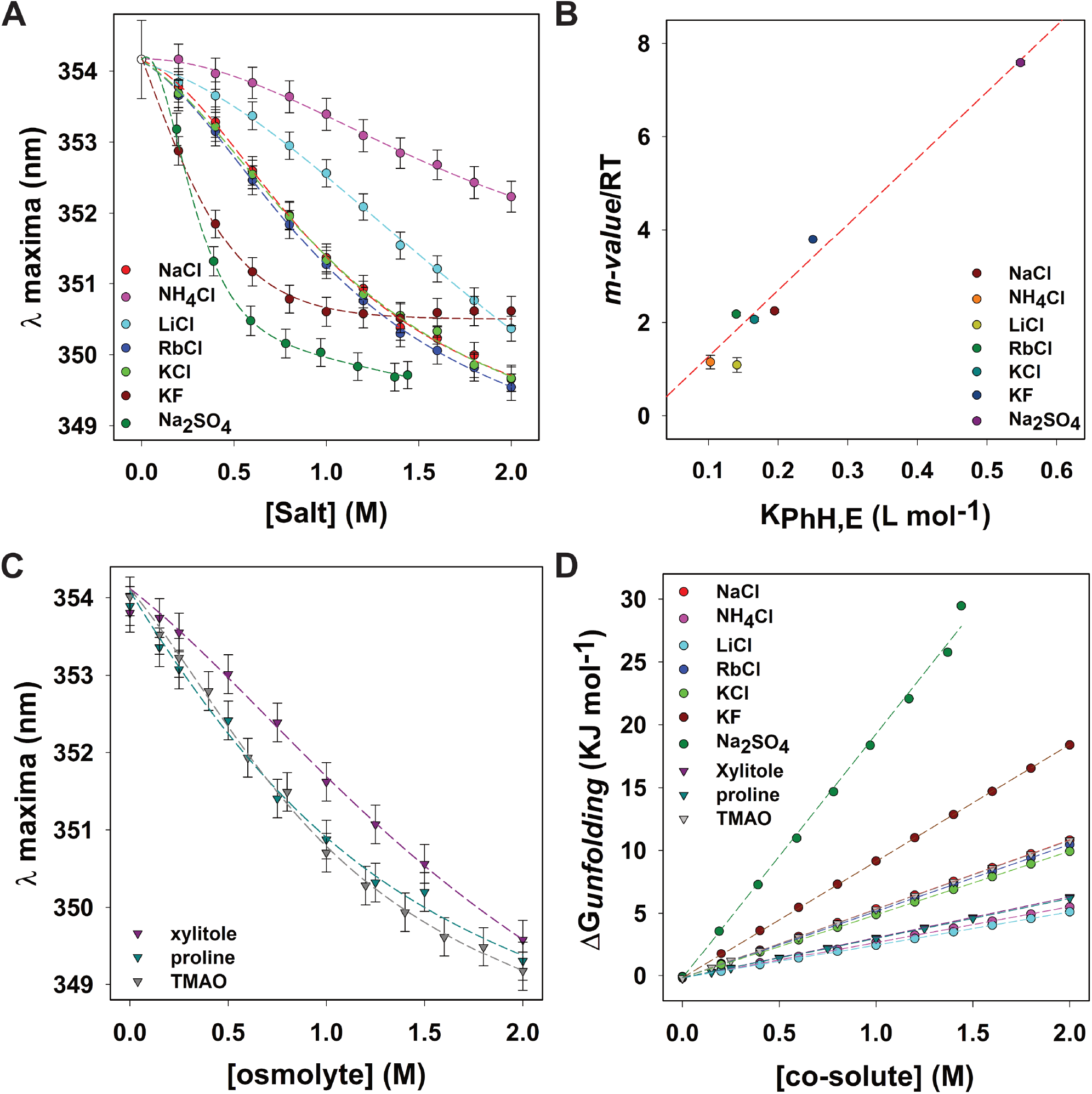
Salt specific effects on the folding thermodynamics of Prx represents hydrophobic collapse. **(A)** Salt and **(B)** osmolyte induced effect on the tertiary structure of PrxF31W as monitored by the intrinsic tryptophan fluorescence emission peaks (λ_maxima_) in various co-solute solutions. All samples were excited at 280 nM and the emission spectra were collected from 300 to 450 nm, with excitation and emission slits both set to 2 nM bandpass. All measurements were performed in triplicate and averaged, the *λ*_maxima_ was obtained by fitting each emission peak to a Weibull 5-parameter regression, the error was obtained from the standard deviation of triplicate measurements. The data were globally fit to Eq-4, constraining *λ*_U_ (the emission maximum of the unfolded protein) to 357.6 nm, while sharing *λ*_F_ (350.5 ± 0.3 nm; the emission maximum of the folded protein) through non-linear least square regression analysis. **(C)** Dependence of 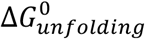 of PrxF31W as a function of co-solute concentration. The data were obtained using the 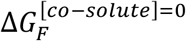 and the “*m-values*” from the global fits of the data depicted in in Figure 7. The dashed lines represent 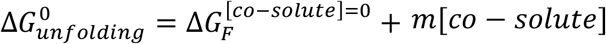 as defined by the parameters tabulated in Table-2. **(D)** Plotting the “*m-value*” parameters from Table 2 against the salting out efficiency as represented by the experimentally determined Setchenow constants obtained from reference 44. The dashed line represents the best linear fit obtained from the data.

We can thus rank the efficiency of each salt in promoting the folded state of Prx, based upon *m*_*salt*_ as:

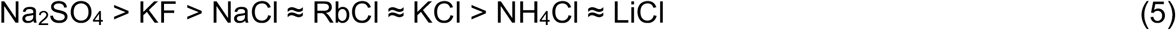

Wohl et al.^43^, have shown that salt-induced conformational changes observed in intrinsically disordered proteins can be caused by salting-out of the hydrophobic residues, provided that the IDP: a) is moderately charged (less than 40 percent of sequence residues), b) has significant hydrophobic content, and c) has polyampholyte rather than polyelectrolyte character (i.e., positive and negative charges balance out each other). At pH 7, Prx has 28 hydrophobic, 14 acidic, and 8 basic residues out of a total of 60; thus, making it a moderately charged polyampholyte with significant hydrophobic content. Therefore, the salt specific effects we observe in the Prx protein folding free energies should be caused by differences in hydrophobic group salting-out efficiencies^40-41^. How efficiently a salt species reduce the solubility of a given hydrophobic molecule is characterized by the Setschenow constant *k* _*set*_ defined as^44^:

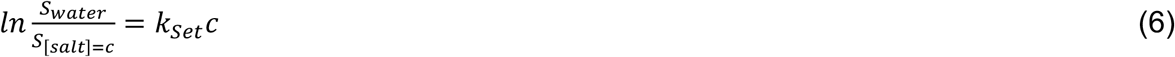

Where *S*_*water*_ is the solubility of the molecule in pure water and *S*_[*salt*]=*c*_ is the solubility of the same molecule in a saline solution having a concentration of [*salt*] = *c*. In Figure 7B, we have plotted our measured *m*_*salt*_ values associated for each salt, compared to its Setschenow constant^45^ for benzene^44^. It can be seen that these values are linearly correlated with each other, confirming that the salt-induced folding of Prx is essentially caused by the salting-out of hydrophobic moieties^41^.

In addition to salts, osmolytes can also induce Prx to fold, as seen in Figure 7C. The data in Figure 7C can be fit to Eq-4a, in which all parameters are defined similar to Eq-4, with osmolyte concentration being substituted for salt:

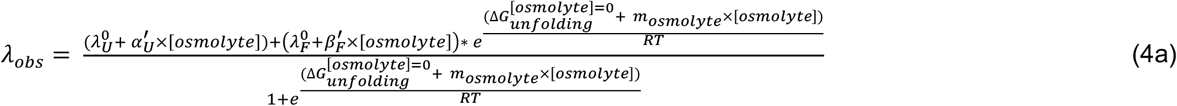

The resulting fitting parameters are tabulated in Table 2. When the concentration dependences of Δ*G*_*unfolding*_ upon salt or osmolyte concentration are plotted in Figure 7D, it becomes clear that they all extrapolate within error to essentially identical values of 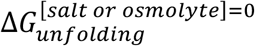. Unlike the salts in Eq-5, which mainly promote protein folding through reducing the solubility of hydrophobic residues, osmolytes stabilize the folded form through reducing the solubility of the polypeptide backbone^46-47^. The fact that the standard Prx unfolding free energy extrapolates to the same value irrespective of perturbing agent, confirms the fact that the conformational itinerary of Prx can be represented by a two-state protein folding model.

## DISCUSSION

Under dilute solvent conditions, Prx exists as a disordered and highly dynamic protein. Additionally, at standard physiological buffer conditions Prx remains disordered. For example, both our CD and NMR measurements show that Prx is unfolded in buffers at pH 7.0 and up to 250 mM salt (Figure 2 and Figure 3). Moreover, both our titration data strongly suggests that Prx does not adopt the fold observed in the X-ray crystal structures until it binds ComR (Figure 3). The CD and NMR data presented here, in addition to our past AUC^14^ and SEC-SAXS^15^ data, clearly demonstrate that Prx is an intrinsically disordered protein (IDP) that only adopts a rigid globular fold when bound to ComR.

The addition of salt or osmolyte causes Prx to fold into a solute-stabilized from via a two-state mechanism (Figure 5, Figure 6, and Figure 7). When we consider how and why salts have a significant effect on the fold of Prx, it is important to note that Prx is an acidic protein and at neutral and physiological pH has a significant excess of negative charges^14^. However, our work demonstrates that the folded form of Prx observed at high salt concentration is not stabilized through the screening of repulsive charges. In fact, salt addition stabilizes a folded form of the protein through salting-out interactions. This result is consistent with the calculations of Wohl et al.,^43^ indicating that salting-out effects play a much more important role when compared to electrostatic screening. Specifically, this is true in salt-induced conformational changes for disordered proteins having a moderate number of charged residues (less than 40 percent). Additionally, the X-ray crystal structures of Prx show that Prx can adopt a highly negatively charged surface. This surface is made of conserved residues that are critical for its interaction with the DBD of ComR. Furthermore, the DBD is highly positively charged and the complementing electrostatics contribute significantly to the nanomolar binding affinity of Prx for ComR^15^. Given the known X-ray structures of Prx^14-15^, that Prx is an IDP (Figure 2 and Figure 3), and our folding analysis (Figure 5, Figure 6, and Figure 7), we now suggest that the Prx:ComR binding mechanism may involve the following equilibria:

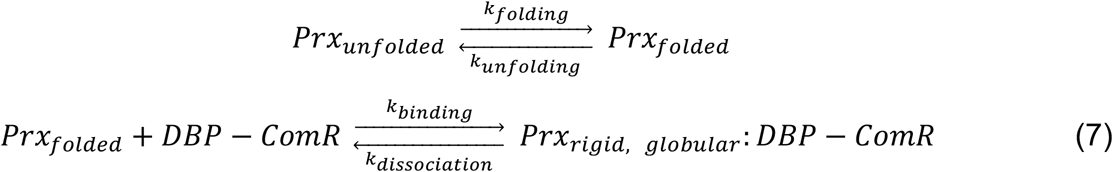

In other words, the ComR-DBD has “salted-out” the rigid globular fold of Prx.

Our salt and osmolyte data also explain how we were previously able to capture a crystal structure of Prx without a binding partner. Typically, IDPs will not crystallize as the process of crystallization requires a high degree of protein order to form a repeating lattice. However, we had purified the C-terminal 6His-tag Prx construct which we show adopts a similar fold to salt-stabilized tag-less Prx (Figure 2 and Figure 3). Also, the crystallization conditions included both salts (0.1 M ammonium acetate and 0.1 M sodium citrate) and a high concentration of crowding osmolytes (30% PEG 4000)^14^. As such, the crystallization conditions and 6His-tag pushed the equilibrium of Prx towards a folded form, and ultimately the rigid globular fold that Prx adopts when salted out by the ComR DBD. Our results also strongly advise that when considering the folding and dynamics of any protein, one must consider the effects of His-tags and other cloning artifacts^29^.

The fact that osmolyte addition stabilizes the folded form of Prx hints to its putative dominant form *in vivo* before binding ComR. Prokaryotic cells are known to be highly crowded with different solutes, RNA, and protein^48^. It is therefore likely that Prx is predominantly in the salt and osmolyte induced folded form when it is first expressed in the GAS cell. However, our NMR data demonstrate that the solute-induced fold is different than the ComR bound fold (Figure 3). Given this, we propose that the Prx binding mechanism of ComR could be described by at least a three-state process, as depicted in Eq-7. First, Prx is in equilibrium between an unfolded state (*Prx*_*unfolded*_), and a solute-stabilized folded state (*Prx*_*unfolded*_) as demonstrated by our data. The high concentrations of crowders and osmolytes existing in the bacterial cell will shift this equilibrium towards *Prx*_*folded*_state. Next, upon encountering ComR, *Prx folded* binds the DBD and adopts a final rigid globular fold by through strong favorable interactions with the highly positive DNA binding surface of the ComR DBD.

The fact that Prx is an IDP and conformationally dynamic may help explain the unknown process of how Prx influences the conformation of ComR. The inactive apo-form of ComR adopts a conformation where the positive DNA binding surface of the DBD is shielded from the solvent and bound tightly to the ComR TPR (tetratricopeptide repeat) domain^20^. This conformation prevents ComR from interacting with DNA until XIP binds the TPR to induce a conformational change that releases the DBD^21^. Like XIP, Prx also changes the conformation of ComR to release the DBD. However, unlike XIP Prx binds the DNA-binding residues of the DBD that are shielded by the TPR^15^. Thus, Prx binding to the apo-conformation of ComR would result in a large steric clash making it unclear how Prx accesses the DBD. Given that Prx can adopt folds other than the rigid form observed in the crystal structures, it is tempting to speculate that the solvent stabilized Prx fold is an intermediate that can first bind ComR and encourage the initial release of the DBD. This would suggest that Prx binds ComR in large part by conformational selection^49-50^. In other words, the solute-stabilized fold (*Prx*_*folded*_) we have observed in this study may be a pre-folded intermediate in the conformational ensemble of Prx that selects for ComR. However, as *Prx*_*folded*_ is not identical to the ComR bound fold of Prx (*Prx*_*rigid, globular*_, Figure 3), the binding mode of Prx to ComR (i.e., *k*_*binding*_), also likely includes an induced-fit component^49-50^. In this case, binding to ComR induces Prx to adopt its final rigid conformation observed in our X-ray co-complex structures^15^.

As we have shown that Prx has a solute-stabilized intermediate form and our NMR data strongly suggests that DBD binding induces additional structure and rigidity within the Prx fold, we hypothesize that Prx is an IDP that binds ComR by a combination of conformational selection and induced fit. However, to fully prove this model will require additional experimentation. For example, studies have shown that IDPs can be forced to switch between conformational selection and induced fit^49, 51^, which may allow us to further probe Prx:ComR binding dynamics.

Research has shown that IDPs play an increasingly important role in the biology and biochemical pathways of bacteria. For example, the protein Bd0108 is an IDP that regulates pilus secretion and the decision of when to eat other bacteria for *Bdellovibrio bacterivorous*^32^, a triplet of small IDPs was found to affect flagellum gene expression and motility in Salmonella^52^, and PopZ (polar organizing protein) is an IDP that forms a protein recruitment hub at bacterial poles^53-54^. Our work here now adds Prx to the short but growing list of known bacterial IDPs. Additionally, in eukaryotes it has been accepted that IDPs and protein IDRs (intrinsically disordered regions) form condensates to serve as “membrane-less organelles” to nucleate protein complex formation and drive biochemical reactions^55^. For example, IDRs in the protein TPX2 create phase-separation condensates essential for microtubule formation^56^. Given the prevalence of IDRs and IDPs in bacteria the same condensate hypothesis for IDP biochemical function has been proposed^57^. As Prx is an IDP this raises the question if it can also form condensates for a particular biochemical function within *Streptococcus*. This question is especially relevant as small phage proteins are typically evolved to perform multiple functions given the small size of a viral genome^58^. As we have previously noted that Prx is functionally similar to the multi-functional phage protein Aqs1 in *Pseudomonas* aeruginosa^15^, we now hypothesize that the IDP nature of Prx may help facilitate interactions with other yet to be discovered binding partners. Overall, the phage protein paratox is a highly dynamic protein whose fold is readily influence by its local environment and protein binding partners.

## MATERIALS AND METHODS

### Protein expression and purification

Expression constructs for Prx included C-terminal 6His-tagged Prx and N-terminal GST-tagged Prx. The 6His-tag Prx construct used was from previous studies^15^. For the N-terminal GST-tagged fusion construct paratox from MGAS315 was placed in the vector pGEX6P1 using BamHI and XhoI sites and ordered from Genscript. For PrxF31W, residue F31 was mutated to W31 using Q5 mutagenesis (New England Biolabs) only in the GST-tagged construct. Expression constructs for ComR included full-length ComR and the minimal ComR DNA-binding domain (DBD) both from *Streptococcus mutans* created in previous studies^14-15^.

Wild-type paratox with a C-terminal 6His-tag from MGAS315 was purified as described previously by nickel chelating affinity chromatography followed by size-exclusion chromatography (SEC)^14-15^. Both full-length ComR and the ComR DBD were also purified as previously described using nickel chelating affinity chromatography followed by size-exclusion chromatography (SEC)^14-15^. The final buffer in SEC was 50 mM Tris pH 7.5 100 mM NaCl 1 mM beta-mercaptoethanol (βME).

GST-tagged Prx proteins were purified as described here. Cells were grown in LB at 37°C to an optical density of 0.8 at 600 nm at which point isopropyl *β*-D1-thiogalactopyranoside IPTG was added to 1mM and the temperature reduced to 20°C. The cells were allowed to grow over-night and then collected by centrifugation and resuspended in lysis buffer (50 mM Tris pH 7.5 250 mM NaCl 2mM beta-mercaptoethanol or *β*ME). Cells were lysed using an Emulsiflex C3 (Avestin) after the addition of 1 mM PMSF, 10 mM MgCl_2_ and DNase I. Lysates were cleared of insoluble debris by centrifugation at 16,000 rpm or 22,000xG. The soluble lysate was then passed over a Q-sepharose gravity column equilibrated in lysis buffer to remove DNA and other contaminants. The flow through was then further purified using a glutathione-Sepharose 4B (Cytiva) gravity column, washed with 500 mL of lysis buffer and then eluted with lysis buffer containing 50 mM glutathione. The GST-tag was removed by the addition of HRV-3C protease with overnight incubation and dialysis into lysis buffer at 4°C. After digestion the sample was passed back over the glutathione-Sepharose 4B gravity column to remove GST and undigested material. Following GST removal, free Prx was further purified by SEC using a HiLoad 16/600 superdex 75 gel filtration column (Cytiva). The final buffer was 50 mM Tris pH 7.5, 100 mM NaCl, and 1 mM *β*ME. Prx samples were concentrated and flash-frozen in liquid nitrogen for later use.

For isotopically labeled Prx protein samples, cells were grown in minimal media (M9) but with the nitrogen source substituted to ^15^N ammonium chloride (Cambridge Isotopes). The subsequent steps of cell growth, lysis, and purification are the same as the unlabeled protein samples.

### Circular dichroism spectroscopy

Wildtype Prx and residue point-variant PrxF31W were dialyzed overnight at 4°C in 1.5 L of CD buffer (20 mM sodium phosphate, pH 7.0) using dialysis membrane with a 2 kDa pore size cut-off. The final protein concentration was determined using a Thermo Scientific™ NanoDrop™ One Microvolume UV-Vis Spectrophotometer (PrxWT abs 0.1% = 0.602; PrxF31W abs 0.1% = 1.336). The protein was then diluted to the appropriate stock concentration using CD buffer. A series of 300 μL samples with a final concentration of 20 μM Prx (WT and F31W) were prepared at various concentrations of potassium fluoride. All CD measurement were performed using a Jasco J-810 spectropolarimeter (Easton, MD). The samples were loaded into a 1 mm pathlength cylindrical cuvette. The spectra were collected in triplicate from 260 nm to 190 nm for each sample and averaged. Xylitol, TMAO, proline, and urea (ultragrade) were purchased from Sigma Aldrich. All salts and dialysis tubing were purchased from Fisher Scientific (Fair Lawn, NJ).

### Nuclear magnetic resonance spectroscopy

^1^H-^15^N HSQC spectra were recorded at 20°C on a Varian Unity Inova 600 MHz spectrometer equipped with room-temperature Varian 5 mm Triple-resonance H/C/N inverse-detection solution probe with Z gradient probe. Samples were prepared for in their various buffers and included 10% D2O. Prx was used at 400 μM concentration in the following buffers all at pH 7.0, 20 mM sodium phosphate, 20 mM sodium phosphate containing 6.66 M urea, 20 mM sodium phosphate containing Na2SO4 (at 200 mM, 500 mM and 700mM). For the Prx and ComR DBD complex, 500 μM of unlabeled DBD was added to 15N labeled Prx in 20 mM sodium phosphate pH 7.0 Data was processed using NMRpipe^59^ and analyzed using UCSF Sparky.

### Fluorescence spectroscopy

Concentrated PrxF31W was dialyzed in 1 L of buffer (25mM Tris pH 7.0) for 1.5 hours to ensure adequate buffer exchange. The protein was then diluted to the appropriate stock concentration. Steady-state fluorescence spectra were measured on a Fluorolog-3 Horiba Jobin Yvon spectrofluorometer (Edison, NJ) using a 10 × 3.3 mm quartz cuvette to hold the sample. The samples were excited at 280 nm, the excitation and emission slits were set to a 2 nm bandpass. All equilibrium folding and unfolding experiments in this work was performed upon protein samples with a final concentration of 5μM PrxF31W in the buffer of choice. All samples were equilibrated at room temperature (20 °C) for 1 hour prior to the scan. In the unfolding experiments, Urea stock solutions were prepared for each predetermined concentration of potassium fluoride. The final urea concentrations were determined by refractive index^60^ using 115V AC/DC Refractometer purchased from Fisher Scientific (Free Lawn, NJ) to verify the final concentration of denaturant post data acquisition. The data were analyzed with Sigma Plot (Point Richmond, CA) software.

### Stopped-flow fluorescence kinetics

Unfolding and refolding kinetics were performed using Applied Photophysics SX-20 (Surrey, UK) stopped-flow fluorescence instrument (dead time ∼ 1 ms). The excitation wavelength was set to 280nm, and the emission was monitored using a 330 nm Bandpass filter (FWHM 10nm). To ensure the appropriate dilution of denaturant, asymmetric mixing was set up using 2.5 ml and 0.25 ml drive syringes purchased from Delta photonics (Ottawa, CA). In the refolding experiments, concentrated PrxF31W was disolved in various urea containing KF solutions, and subsequently diluted 10-fold into the appropriate salt buffer upon mixing. The unfolding experiments were performed in a similar manner whereby, the protein was folded in either 0.5 M or 1.0 M KF solutions, and then diluted 10-fold into various urea containing KF solutions. The final protein concentration was between 1-2 μM and the final urea concentration was determined by refractive index post mixing. All kinetic experiments were done at 20 °C and each measurement was performed at least 15 times and averaged. The kinetic parameters were determined by fitting the data to a mono-exponential equation using Sigma Plot (Point Richmond, CA) software.

## ACKNOWLEDGEMENTS

We would like to thank Tasneem Hassan Muna in the Prehna lab for guidance and training with protein purification.

## FUNDING AND ADDITIONAL INFORMATION

This work was supported by the Natural Sciences and Engineering Research Council of Canada (NSERC) RGPIN-2018-04968 to G.P., and RGPIN-2017-05935 to M.K., and a Canadian Foundation for Innovation award (CFI) 37841 to G.P., and 23175 to M.K.; I.A. was the recipient of a University of Manitoba Graduate Fellowship.

